# A consensus protocol for the recovery of mercury methylation genes from metagenomes

**DOI:** 10.1101/2022.03.14.484253

**Authors:** Eric Capo, Benjamin D. Peterson, Minjae Kim, Daniel S. Jones, Silvia G. Acinas, Marc Amyot, Stefan Bertilsson, Erik Björn, Moritz Buck, Claudia Cosio, Dwayne Elias, Cynthia Gilmour, Maria Soledad Goñi Urriza, Baohua Gu, Heyu Lin, Yu-Rong Liu, Katherine McMahon, John W. Moreau, Jarone Pinhassi, Mircea Podar, Fernando Puente-Sánchez, Pablo Sánchez, Veronika Storck, Yuya Tada, Adrien Vigneron, David Walsh, Marine Vandewalle-Capo, Andrea G. Bravo, Caitlin Gionfriddo

**Affiliations:** Department of Marine Biology and Oceanography, Institute of Marine Sciences, CSIC, Barcelona, 08003, Spain; Department of Aquatic Sciences and Assessment, Swedish University of Agricultural Sciences, Uppsala, 75007, Sweden; Department of Bacteriology, University of Wisconsin at Madison, Madison, WI 53706, United States; Natural Resource Ecology Laboratory, Colorado State University, Fort Collins, CO 80523, USA; Department of Earth and Environmental Science, New Mexico Institute of Mining and Technology, Socorro, NM 87801, USA; National Cave and Karst Research Institute, Carlsbad, NM 88220, USA; Department of Biological Sciences, University of Montréal, Montréal, QC, H3C 5J9, Canada; Department of Chemistry, Umeå University, Umeå, 90736, Sweden; University of Reims Champagne-Ardenne, UMR-I 02 SEBIO, Reims, 51100, France; Oak Ridge National Lab, Oak Ridge, TN 37830, USA; Smithsonian Environmental Research Center, Edgewater, MD 21037, USA; University of Pau et des Pays de l’Adour, E2S UPPA, CNRS, IPREM, Pau, 64000, France; School of Geography, Earth and Atmospheric Sciences, The University of Melbourne, Parkville, VIC 3010, Australia; College of Resources and Environment, Huazhong Agricultural University, Wuhan, 430070, China; School of Geographical and Earth Sciences, University of Glasgow, Glasgow, G12 8RZ, UK; Centre for Ecology and Evolution in Microbial Model Systems - EEMiS, Linnaeus University, Kalmar, 39231, Sweden; National Institute for Minamata Disease, Department of Environment and Public Health, Kumamoto, 867-0008, Japan; Department of Biology, Concordia University, Montreal, Quebec H4BIR6, Canada

**Keywords:** mercury, *hgcAB* genes, Hg methylation, metagenomics, bioinformatics, Hg-MATE, marky-coco

## Abstract

Mercury methylation genes (*hgcAB)* mediate the formation of the toxic methylmercury and have been identified from diverse environments, including freshwater and marine ecosystems, Arctic permafrost, forest and paddy soils, coal-ash amended sediments, chlor-alkali plants discharges and geothermal springs. Here we present the first attempt at a standardized protocol for the detection, identification and quantification of *hgc* genes from metagenomes. Our Hg-MATE (Hg-cycling Microorganisms in Aquatic and Terrestrial Ecosystems) database, a catalogue of *hgc* genes, provides the most accurate information to date on the taxonomic identity and functional/metabolic attributes of microorganisms responsible for Hg methylation in the environment. Furthermore, we introduce “marky-coco”, a ready-to-use bioinformatic pipeline based on *de novo* single-metagenome assembly, for easy and accurate characterization of *hgc* genes from environmental samples. We compared the recovery of *hgc* genes from environmental metagenomes using the marky-coco pipeline with an approach based on co-assembly of multiple metagenomes. Our data show similar efficiency in both approaches for most environments except those with high diversity (i.e., paddy soils) for which a co-assembly approach was preferred. Finally, we discuss the definition of true *hgc* genes and methods to normalize *hgc* gene counts from metagenomes.

## Introduction

Environmental mercury methylation is primarily a biotic process carried out by microorganisms that transform inorganic mercury (Hg) into the more toxic and bioaccumulative monomethylmercury (MeHg). The capacity to perform Hg methylation was historically associated with certain sulfate-reducing bacteria, iron-reducing bacteria and methanogenic archaea (Compeau and Bartha, 1985; Fleming et al., 2006; Kerin et al., 2006; Hamelin et al., 2011). Field observations revealed links between Hg methylation and sulfate-reduction, iron-reduction and methanogenesis in organic matter-rich anaerobic environments (Bravo and Cosio, 2020 for review), as well as subsequent studies that tested cultured representatives of these clades for Hg-methylation capability (Fleming et al. 2006; Gilmour et al., 2011; 2013; 2018). The discovery of the *hgc* genes (Parks et al., 2013) has facilitated the detection of novel putative Hg methylating bacteria and archaea through cultivation-independent molecular methods (Podar et al., 2015; Gionfriddo et al., 2016). Recent works analyzing publicly available genomes and environmental metagenome-assembled genomes (MAGs) identified *hgc*-containing *(hgc*^+^) microorganisms from microbial lineages not formerly associated with Hg-methylation, such as members of the PVC superphylum (Jones et al., 2019; Gionfriddo et al., 2019; Peterson et al., 2020; McDaniel et al., 2020; Lin et al., 2021). Identifying *hgc* genes in microbial genomes from meta-omic datasets greatly expanded our view of the phylogenetic diversity of putative Hg methylators (Fig 1), but we still do not fully understand which microorganisms are the main drivers of Hg methylation in diverse environments, particularly outside of anoxic sediments.

**Figure 1.**
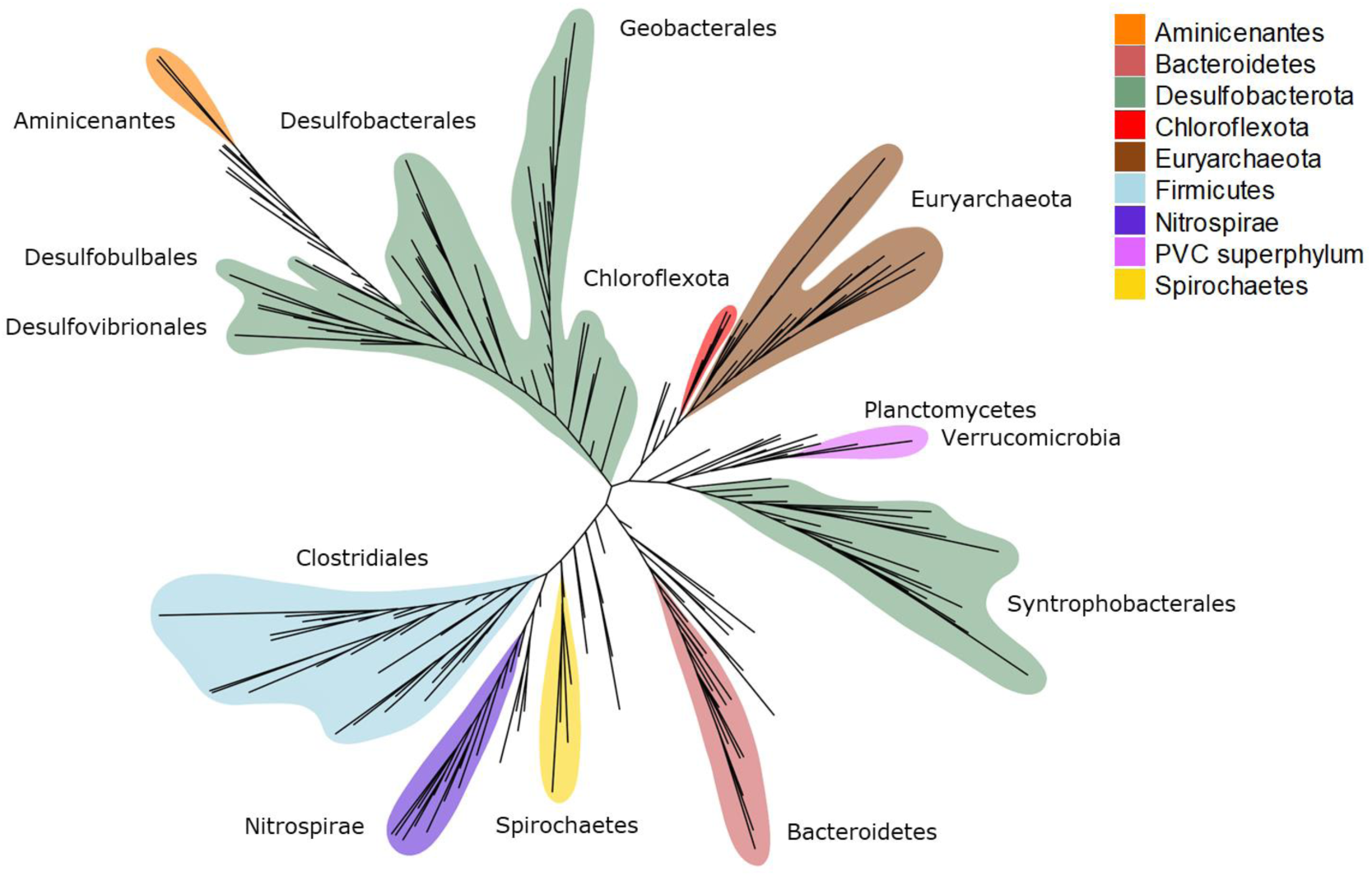
Simplified unrooted phylogenetic tree of *hgcA* sequences from the Hg-MATE database. Taxonomy is based on NCBI classification with the exception of Deltaproteobacteria (Desulfobacterota with GTDB classification) and Chloroflexi (Chloroflexota with GTDB classification). For visualization ease, microbial groups were collapsed by the dominant monophyletic group. Microbial groups with the highest diversity of *hgc*^+^ microorganisms are denoted by colors.

Significant knowledge gaps in the identification of microorganisms capable of Hg methylation remain, largely because of the absence of *hgc*^+^ cultured representatives from novel clades (i.e., outside the Desulfobacterota, Firmicutes, Methanomicrobia) with experimentally validated Hg-methylating capability (Gilmour et al., 2018). One reason for this is the difficulty in selecting for *hgc*^+^ microorganisms during cultivation, and another is the lack of a successful methodology for isolating all relevant microbes in controlled laboratory conditions. Microbes that have yet to be cultivated, and for which successful laboratory growth parameters need to be identified, are often referred to as the “unculturable” (Hug et al., 2016; Steen et al., 2019). High-throughput meta-omic and targeted amplicon sequencing studies have become the main methods for identifying putative Hg methylating microorganisms of this unculturable fraction (Bravo et al., 2018; Gionfriddo et al., 2020; Xu et al., 2021). While directly testing for Hg methylation capacity may not be a viable strategy, pairing these sequencing methods with biogeochemical measurements, Hg methylation assays, and other manipulation studies can connect a Hg-methylating microbiome to MeHg production and metabolic activity and help to elucidate the potential contribution of these novel clades to Hg methylation (Kronberg et al., 2016; Bouchet et al., 2018; Schaefer et al., 2020; Roth et al., 2021).

The detection of hgc^+^ MAGs provides the most precise information about the taxonomic and metabolic characteristics of putative Hg methylators (Jones et al., 2019; Peterson et al., 2020; Lin et al., 2021; Vigneron et al., 2021). However, the microbial diversity in some environments is too high and/or Hg methylators are too rare to identify them effectively (Podar et al., 2015; Christensen et al., 2019). In these cases, read-based metagenomic analyses and *hgc* metabarcoding are easier and more economical. Accurately identifying Hg-methylating clades (and metabolic guilds) from *hgc* sequences alone therefore requires a universally used and updated *hgcAB* database, coupled to consistent and robust bioinformatic practices, in order to identify precisely the target genes in complex meta-omic datasets.

In this work, we introduce Hg-MATE (Hg-cycling Microorganisms in Aquatic and Terrestrial Ecosystems) database version 1 (https://doi.org/10.25573/serc.13105370.v1), an up-to-date *hgcAB* catalog compiled from isolated, single-cell and metagenome-reconstructed genomes. Additionally, we present marky-coco (https://github.com/ericcapo/marky-coco), a ready-to-use bioinformatic pipeline to detect, identify and count *hgc* genes from metagenomes (**Fig 2**). We apply this pipeline to metagenomes collected from paddy soils, brackish and lake waters, as well as sediments from reservoirs and lakes in which *hgc* genes have been previously detected (Liu et al., 2018; Jones et al., 2019; Capo et al., 2020; Millera Ferriz et al., 2021). Further, we specifically compared the reliability of (i) applying the marky-coco pipeline based on *de novo* single assembly approach from single metagenomes with (ii) co-assembling of multiple metagenomes (co-assembly) prior to mapping and identification. Finally, we discuss appropriate definitions and cutoff criteria for *hgc* genes and also best practices to normalize data for an accurate count of *hgc* genes in metagenomes from environmental samples.

**Figure 2.**
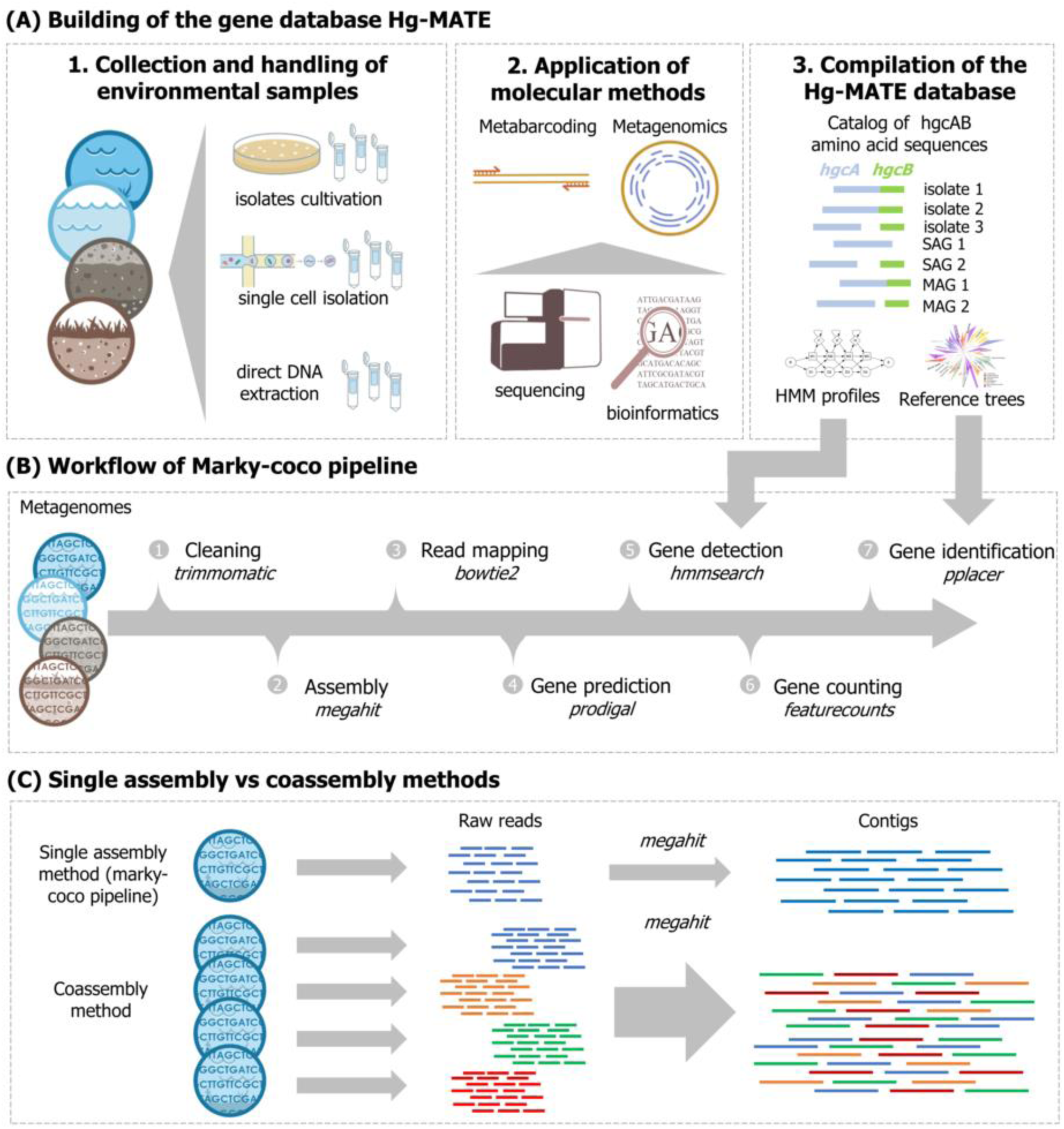
(A) Workflow illustrating how the *hgcAB* gene catalogue Hg-MATE database was built, (B) Simplified workflow of the marky-coco pipeline (C) Illustration of the two assembly approaches compared in this work: single assembly vs co-assembly.

## 2. Material and Methods

### 2.1 Description of the Hg-MATE database v1

The Hg-MATE database v1 was released on 14 January 2021 (https://doi.org/10.25573/serc.13105370.v1), and contains an extensive *hgcAB* dataset from a wide range of microorganisms and environments. The catalog contains 1053 unique HgcA/B amino acid sequences (Table 1). We categorized the HgcAB amino acid sequences into four types depending on whether they were encoded in (i) pure culture/environmental microbial isolates (ISO) (ii) single-cell genome sequences (CEL) (iii) metagenome-assembled genomes (MAGs) (iv) or an environmental meta-omic contig (CON). Amino acid sequences of HgcA, HgcB, and concatenated HgcA and HgcB were included in the database. If *hgcB* was not co-localized with *hgcA* in the genome and/or could not be identified, then ‘na’ was listed in the ‘HgcB’ sequence column. Both genes need to be present and encode functional proteins for a microbe to methylate Hg (see Parks et al., 2013; Smith et al., 2015). One reason *hgcB* may not be identified in some genomes carrying *hgcA* is because HgcB is highly homologous to other 4Fe-4S ferredoxins. Therefore, *hgcB* can be difficult to differentiate from other ferredoxin-encoding genes if not co-localized with *hgcA* on a contiguous sequence. In addition, *hgcB* may be missing from ‘MAGs’, ‘CEL’ and ‘CON’ sequences due to incomplete coverage of the genome or incomplete contig assembly, or failure to bin the contig carrying *hgcB*. Some *hgc* genes are predicted to encode a ‘fused HgcAB protein’ which has been previously described (Podar et al., 2015), and is characterized by one gene that encodes for a 4Fe-4S ferredoxin-like protein with shared homology to HgcA and HgcB. This ‘fused HgcAB’ protein contains the corrinoid iron-sulfur and transmembrane domains characteristic of HgcA as well as the 4Fe-4S ferredoxin motif of HgcB (e.g., Uniprot Q8U2U9, NCBI Refseq: WP_011011854.1, *Pyrococcus furiosus* DSM 3638). These sequences are provided in the ‘HgcA’ column, and labeled ‘fused HgcAB’ in the HgcB column. These ‘fused HgcAB’ sequences should be treated with caution because, while they share significant sequence homology to HgcA and HgcB from confirmed Hg methylators, to date all organisms with a ‘fused HgcAB’ that have been tested do not seem to produce MeHg in culture (Podar et al., 2015; Gilmour et al., 2018).

**Table 1.** Summary of HgcAB sequence types in version 1 of the Hg-MATE database.

| Genome type | Total HgcA(B) sequences | Encode both HgcA and HgcB | Encode fused HgcAB | Only HgcA (or HgcB) present |
| --- | --- | --- | --- | --- |
| ISO | 204 | 173 | 10 | 21 |
| CEL | 29 | 4 | 18 | 7 |
| MAG | 787 | 696 | 17 | 74 |
| CON | 33 | 9 | 0 | 21(3) |

The resources within the Hg-MATE database v1 include a catalog with the amino acid sequences and metadata of all microorganisms. Only sequences with genomic identifying information (i.e., ‘ISO’, ‘CEL’, ‘MAG’) were used to compile further resources. Resources include: (i) FASTA files containing Hgc amino acid sequences; (ii) Multiple Sequence Alignments (MSA) in FASTA format of Hgc amino acid sequences built with MUSCLE implemented in MEGAX (Kumar et al., 2018) with the cluster method UPGMA; and (iii) Hidden Markov models (HMM) of aligned Hgc amino acid sequences built from MSAs using the *hmmbuild* function from the hmmer software (3.2.1 version, Finn et al., 2011). Additionally, resources include reference packages that can be used to identify and classify: (1) the corrinoid-binding domain of HgcA which corresponds to residues ∼37-156 of the HgcA sequence from *Pseudodesulfovibrio mercurii* ND132 and includes the characteristic cap helix domain (2) full HgcA sequence and (3) concatenated HgcA and HgcB. Each reference package contains sequence alignments, an HMM model, a phylogenetic tree, and NCBI taxonomy. Reference packages were constructed using the program Taxtastic (https://github.com/fhcrc/taxtastic) for HgcA(B) amino acid sequences from ISO, CEL & MAG. Phylogenetic trees were built from MSA files by RAxML using the GAMMA model of rate heterogeneity and LG amino acid substitution matrix (Le and Gascuel, 2008). Trees were rooted by HgcA paralog sequences, carbon monoxide dehydrogenases (PF03599) from non-HgcA coding microorganisms Candidatus Omnitrophica bacterium CG1_02_41_171 and *Thermosulfurimonas dismutan*s. These organisms were chosen because of their distant phylogenetic relationship to hgcA^+^ microorganisms. Confidence values on branches were calculated from 100 bootstraps. Using the HgcA reference tree, a simplified tree of ‘ISO’, ‘CEL’, ‘MAG’ *hgcA* genes was built using iTOL (Letunic and Bork, 2019) and clades were collapsed by the dominant monophyletic group, when possible, for visualization ease.

### 2.2 Data collection

A total of 29 metagenomes from recent studies studying *hgc* genes in environments with known active Hg methylation were used for the bioinformatic analyses performed in this work (Table 1, Datasheet 1A). Metagenomes from brackish waters (BARM8s) were collected in 2014 in the Gotland Deep basin of the Central Baltic Sea. Out of 81 available metagenomes (Alneberg et al., 2018; BioProject ID PRJEB22997), 8 metagenomes where *hgc* genes have been detected (Capo et al., 2020) were used in the present analysis. Water depths of these metagenomes ranged from 76 to 200 m with oxygen concentrations either low (hypoxic zone) or undetectable (anoxic zone), salinity ranging between 9.2-12.1 psu and MeHg concentrations measuring up to 1640 fM (Soerensen et al., 2018). Lake sediments and water metagenomes (MANGA6s) were obtained in 2013-2014 from the sulfate-impacted Manganika lake in Northern Minnesota (Jones et al., 2019, BioProject ID PRJNA488162). This hypereutrophic lake is characterized by dissolved oxygen approaching 16 mg/L (nearly 200% saturation) near the surface, pH exceeding 8.7 and MeHg accumulating over 3 ng/L in bottom waters. Dissolved oxygen and pH decreased with depth, and anoxic conditions were encountered below 4 m. Sulfide concentrations up to 2 mM were observed in bottom waters and sediments. Water samples were collected at these anoxic depths. Five metagenomes (RES5S) were obtained from reservoir sediments from the St. Maurice River near Wemotaci, Canada in 2017 and 2018 (Millera-Ferriz et al., 2021, GOLD-JGI Ga0393614 Ga0393582, Ga0393617, Ga0393586, Ga0393589). The studied river section has been affected by the construction of two run-of-river power plant dams and its watershed has been disturbed by a forest fire, logging, and the construction of wetlands. MeHg concentrations in samples varied from <0.02 to 19 ng/g. Metagenomes from paddy and upland soils (PADDY10s) were collected from two historical Hg mining sites, Fenghuang (FH) and Wanshan (WS), in Southwest China in August 2016 (Liu et al., 2018, BioProject ID PRJNA450451). The pH of paddy soils ranged from 6 to 7.5. Historical discharge from Hg mining operations and ongoing atmospheric deposition contribute to high concentrations of MeHg in the soils around these areas with values up to 7.9 ng g^-1^ in the collected samples.

### 2.3 Bioinformatics

The detection, taxonomic identification and counting of *hgc* genes was done with the marky-coco snakemake-implemented pipeline (https://github.com/ericcapo/marky-coco). A brief overview of this workflow is as follows: the metagenomes were trimmed and cleaned using fastp (Chen et al., 2018) with the following parameters: quality threshold of 30 (-q 30), length threshold of 25 (-l 25), and with trimming of adapters and polyG tails enabled (--detect_adapter_for_pe --trim_poly_g --trim_poly_x). A *de novo* single assembly approach, in which each metagenome was assembled individually, was applied using the assembler megahit 1.1.2 (Li et al., 2016) with default settings. The annotation of the contigs for prokaryotic protein-coding gene prediction was done with the software Prodigal 2.6.3 (Hyatt et al., 2010). The DNA reads were mapped against the contigs with bowtie2 (Langdmead and Salzberg, 2012), and the resulting .sam files were converted to .bam files using samtools 1.9 (Li et al., 2009). The .bam files and the prodigal output .gff file were used to estimate read counts by using featureCounts (Liao et al., 2014). In order to detect *hgc* homologs, HMM profiles derived from the Hg-MATE database v1 were applied to the amino acid FASTA file generated with Prodigal from each assembly with the function *hmmsearch* from HMMER 3.2.1 (Finn et al., 2011). The reference package ‘hgcA’ from Hg-MATE.db was used for phylogenetic analysis of the HgcA amino acid sequences. Briefly, the predicted amino acid sequences from gene identified as putative *hgcA* gene were (i) compiled in a FASTA file, (ii) aligned to the Stockholm formatted HgcA alignment from the reference package with the function *hmmalign* from HMMER 3.2.1 (iii) placed onto the HgcA reference tree and classified using the functions *pplacer, rppr* and *guppy_classify* from the program pplacer (Matsen et al., 2010). For more details, see the README.txt of the Hg-MATE database v1 (https://doi.org/10.25573/serc.13105370.v1). Additionally, to compare the efficiency of the marky-coco pipeline to detect *hgc* genes from metagenomes with a co-assembly approach (multiple metagenomes used for assembly), we performed co-assemblies on metagenomes within each environmental system (BARM8s, MANGA6s, RES5s, PADDY10s, Table 2). Post-assembly, all other steps of the analysis procedure were performed similarly to the marky-coco pipeline. Detection of *dsrA* genes were detected in metagenomes with the function *hmmsearch* and HMM profile from TIGRFAM (Selengut et al., 2007). The amount of sequencing required to cover the total diversity and the estimated diversity of each metagenome were evaluated using the Nonpareil method (Rodriguez-R and Konstantinidis, 2014).

**Table 2.** Metagenomes collected from previously published papers investigating the presence of Hg methylators in the environment.

| Systems | #metagenomes | dataset id | References |
| --- | --- | --- | --- |
| Brackish waters | 8 | BARM8s | Capo et al., 2020 |
| Lake sediments and water | 6 | MANGA6s | Jones et al., 2019 |
| River/reservoir sediments | 5 | RES5s | Millera Ferriz et al., 2021 |
| Paddy soils/upland soils | 10 | PADDY10s | Liu et al., 2018 |

### 2.4 Stringency cut-offs for the definition of true *hgc* genes

Based on knowledge from confirmed isolated Hg methylators, we propose several stringency cutoffs that could be used to distinguish between an *hgcA* gene homolog and an *hgcA*-like gene that encodes for a protein of unknown Hg methylation capability. (i) High stringency cutoff: amino acid sequence includes one of the cap-helix motifs with the conserved cysteine (Cys93 in *P. mercurii* ND132), NVWCAAGK, NVWCASGK, NVWCAGGK, NIWCAAGK, NIWCAGGK or NVWCSAGK. This cutoff is based on previous findings that showed isolated microorganisms carrying HgcA proteins with the cap helix domain are capable of Hg methylation (Parks et al., 2013; Smith et al., 2015; Gilmour et al., 2018; Cooper et al., 2020). Within the high stringency cutoff, there is a possible need to distinguish between the amino acid sequences from fused HgcAB-like proteins and those from true HgcA proteins, since isolates that encode fused HgcAB-like genes do not have the capacity to methylate Hg in culture (Podar et al., 2015; Gilmour et al., 2018). The fused HgcAB include the cap-helix and ferredoxin motifs of HgcA and HgcB. (ii) Moderate stringency cutoff: in addition to amino acid sequences that include the motifs described above, any sequence with a bitscore value obtained from the HMM analysis greater than or equal to 100 is included (iii) Low stringency cutoff: in addition to amino acid sequences that include the motifs described above, any sequence with a bitscore value greater than or equal to 60 is included. For *hgcB* gene homologs, we propose two cutoffs that could be used for their description as *hgcB* genes. (i) High stringency cutoff: their amino acid sequences include one of the following motifs featuring the conserved Cys (Cys73 in *P. mercurii* ND132, Cooper et al., 2020), C(M/I)ECGA motifs and that the genes are found on the same contig as an *hgcA* genes. (ii) Moderate stringency cutoff: amino acid sequences include the C(M/I)ECGA motif, but the gene are not co-located on a contig with an *hgcA* gene.

### 2.5 Estimation of *hgcA* abundance in metagenomes

Coverage values of *hgcA* genes were calculated, for each gene and each sample, as the number of reads mapping to the gene divided by the length of the gene (read/bp). We compared the reliability of four procedures for normalizing read counts of *hgcA* genes. Normalization metrics were (i) the total number of mapped reads (ii) the summed coverage values of *rpoB* genes, (iii) the median coverage values of 257 marker genes (GTDB-Tk r89 release, Chaumeil et al., 2019), or (iv) the genome equivalents values calculated using the software MicrobeCensus (Nayfach and Pollard, 2015) which normalizes the relative abundance by the metagenomic dataset size and the community average genome size of the microbial community. The coverage of each marker gene was calculated as the sum of the coverages of all the ORFs assigned to that gene (Datasheet 1A). The *rpoB* and the 256 other marker genes were detected using the function *hmmsearch* from hmmer software (v3.2.1, Finn et al., 2011) and applying the trusted cut-off provided in HMM files (GTDB-Tk r89 release, Chaumeil et al., 2019).

### 2.6 Data analysis

A non-metric multidimensional scaling analysis (nMDS) was performed applying the function *metaMDS* from the R package vegan (Oksanen et al., 2015) to the table of *hgcA* gene coverage values, clustered at the lowest level of NCBI taxonomic identification (txid), obtained with single assembly and co-assembly approaches (Datasheet 1B). A PROTEST permutation procedure analysis (1000 permutations) was performed using the function Procrustes to evaluate the level of concordance of the outputs between both approaches. The functions *rcorr* from the R package Hmisc (Harrell and Harrell, 2019), *corrplot* from the R package corrplot (Taiyun et al., 2017) and *plot3D* from the R package rgkl (Adler et al., 2019) were used to investigate correlations between normalization methods.

## 3. Results

### 3.1 Dataset outputs

A total of 29 single assemblies (one for each metagenome) and 4 co-assemblies (reads from each of the BARM8s, MANGA6s, RES5S, and PADDY10s metagenome sets assembled together) were used to compare the efficiency of a single assembly using the marky-coco pipeline and a co-assembly approach to detect, identify and count *hgc* genes from metagenomes (**Fig. 2**). The number of mapped reads of the analyzed metagenomes ranged between 10.2-110.9 M reads (average, 29.4 ± 19.6) with single assembly and 16.6-120.7 M reads (average, 36.0 ± 19.9) with co-assembly, with the percentage of mapped reads ranging between 16-76 % and 24-89 %, respectively (Datasheet 1A). Nonpareil diversity index values (N_*d*_) of metagenomes were between 18.7 and 23.7 with the highest found in paddy soil metagenomes (Fig. S1, Table 3). Nonpareil curves showed that paddy soil samples from this study required the highest sequencing effort for nearly complete coverage followed by reservoir sediments, and then lake sediment and lake waters and brackish waters (Fig S1). Estimated coverage of paddy soils metagenomes was relatively low (average, 0.30-0.37) compared to other metagenomes (0.49-0.83) showing that only a portion of the diversity of these environmental samples was recovered despite the relatively high sequencing depth (88.6 ± 5.6 M reads) (Table 3). Seven metagenomes (S02, S03, S19, S22, S26, S28, S29) that were used in coassemblies but with low *hgcA* coverage values (i.e., <0.40 obtained from co-assemblies) were not used for further comparison analysis. The remaining 22 metagenomes, labeled MG01 - MG22, had *hgcA* unnormalized coverage values between 0.44 and 3.06 (1.22 ± 0.79) (Datasheet 1A). Only *hgcA* genes (and not *hgcB*) from these metagenomes were used for comparison of the two assembly approaches as *hgcAB* gene pairs were not 100 % similar between the two approaches (Datasheet 1B). Additionally, *hgcAB*-like homologs that are predicted to encode for fused HgcAB proteins were excluded from further analysis.

**Table 3.**
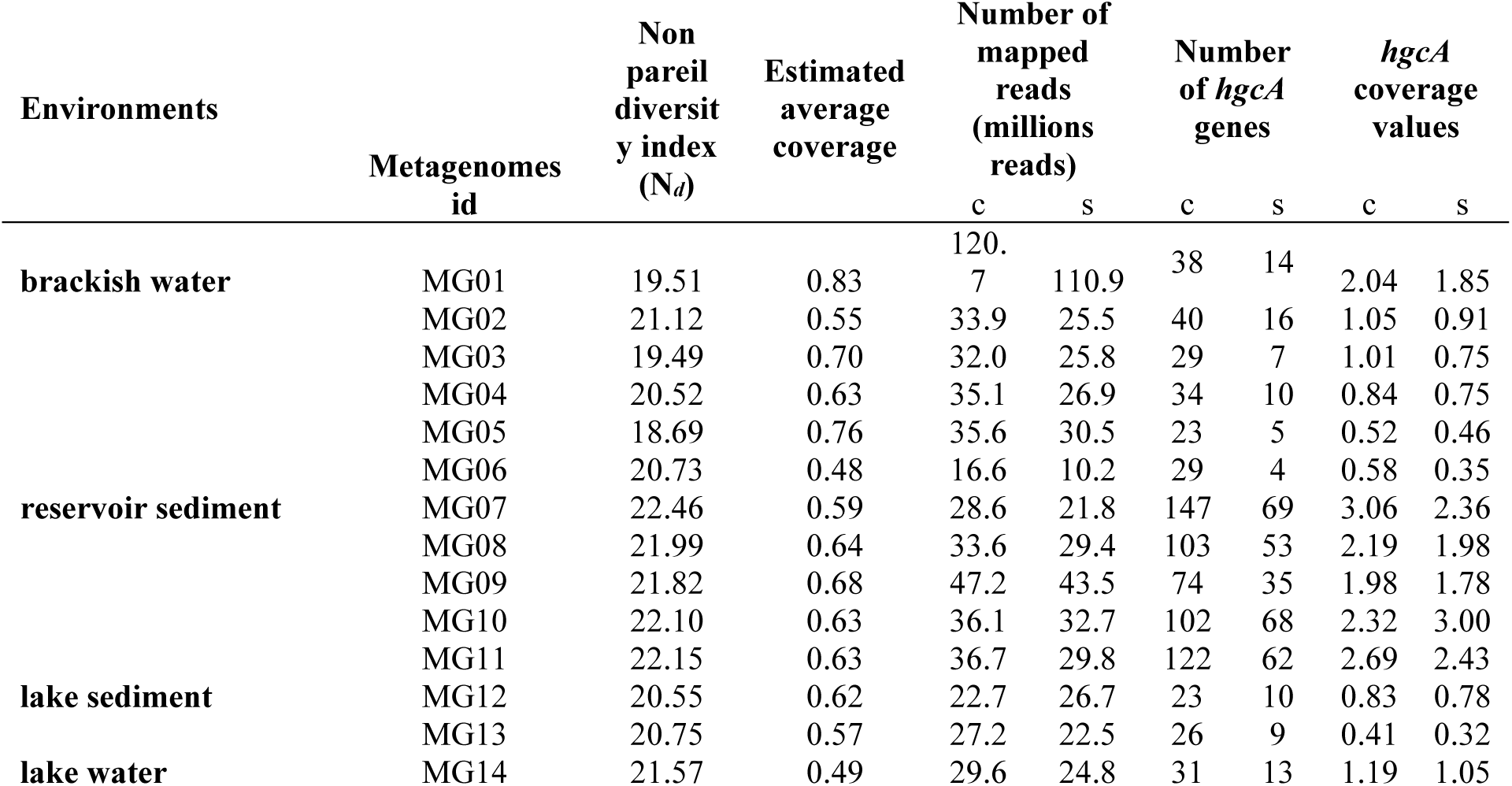
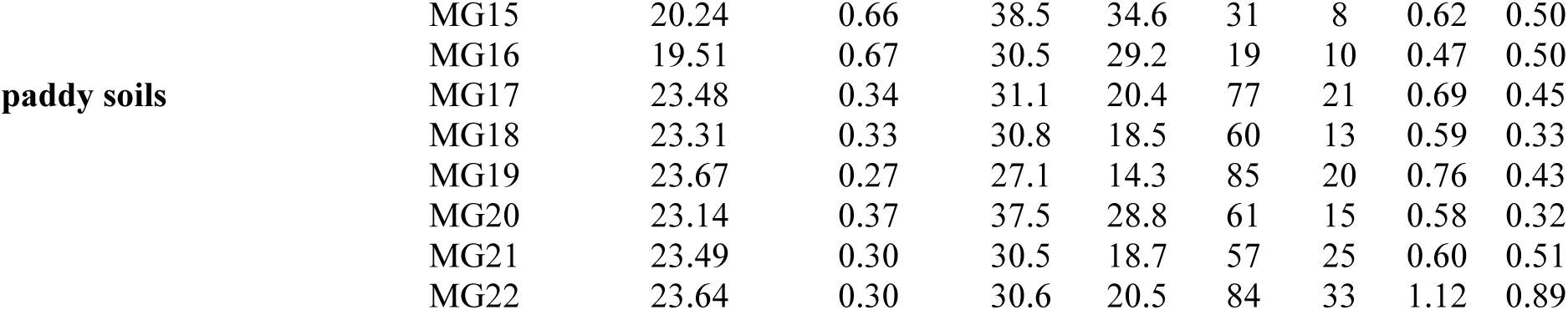
For each metagenome, Non-pareil diversity index values, estimated average coverage, number of mapped reads, number of *hgcA* genes and *hgcA* coverage values (reads/bp) for co-assembly ‘c’ and single assembly ‘s’ approaches. See Datasheet 1A for extended description of the dataset.

### 3.2 Distribution of *hgcA* genes with different stringency cutoffs

By definition, all *hgcA* genes detected with the high stringency cutoff are predicted to encode proteins that include the conserved amino acid motifs characteristic of functional HgcA proteins, while this is not the case for those additionally detected when lowering the stringency cutoffs (i.e., moderate or low). We therefore considered that gene homologs to *hgcA* found with bitscore values below 100 and without conserved motifs cannot with confidence be defined as true *hgcA* genes. Nevertheless, we wanted here to highlight how “false” *hgcA* genes i.e., detected without the conserved amino acid motifs characteristic of functional HgcA proteins, were taxonomically assigned using the *pplacer* approach applied to the Hg-MATE *hgcA* reference tree. The *hgcA* genes detected with a high stringency cutoff and those additionally detected with moderate stringency cutoffs were predominantly identified as Desulfobacterota, Chloroflexota and Euryarchaeota (Fig S2). In contrast, the *hgcA* genes additionally detected with low stringency cutoff were primarily identified as members of the PVC superphylum but were unclassified at lower taxonomic levels. For further comparison, we used information only from *hgcA* genes detected with the high stringency.

### 3.3 Comparison between co-assembly vs single assembly approaches

For all metagenomes, 1.50-7.25 times more *hgcA* genes were detected in co-assemblies (19-147 genes) compared to linked single assemblies (4-69 genes) (Table 3). We investigated the differences in *hgcA* gene lengths, discriminating between genes (i) found at the extremity of contigs (potentially truncated) and (ii) between other genes in contigs therefore expected to be complete. A higher number of ‘complete’ *hgcA* gene sequences were detected with the co-assembly (1-17, average 6.8 ± 4.4 genes) compared to the single assembly (0-6, average 2.0 ± 2.7 genes), e.g., for metagenomes from brackish and lake waters (Datasheet 1A). No complete genes were identified in the single assemblies that were not also identified in the co-assembly. Violin plots illustrated that, overall, a higher number of ‘complete’ *hgcA* sequences (> 950 bp) were found with the co-assembly versus the single assembly (Fig. S3).

In a comparison of HgcA amino acid sequences recovered from the two assembly approaches, no HgcA sequence from the single assembly had 100% sequence identity to sequences in the co-assembly (Datasheet 1B). The highest sequence similarity of HgcA sequences from different assemblies of the same dataset was 99%. To compare, we investigated differences between assemblies for detecting *dsrA* gene, which encodes for dissimilatory sulfite reductase subunit A, an essential enzyme in sulfate reduction and expected to be present in these datasets. Identical amino acid sequences of DsrA-encoding genes were found when comparing single assemblies to the related co-assembly with numbers ranging from 1 to 33 depending on metagenomes (Datasheet 1D). Comparatively, *dsrA* genes were 3-34x more abundant (in coverage) than *hgcA* genes. This higher abundance helps explain why more identical *dsrA* were found between co-assembly and single assembly approaches than for *hgcA* genes.

Distribution plots showed unnormalized coverage values of *hgcA* clustered by environment types (Fig. 3A) or for each metagenome (Fig. S4). Importantly, unnormalized values were used here to compare single assembly vs coassembly results for each metagenome but not to compare difference between environments for which normalization would be required (Fig S4). Overall, higher *hgcA* coverage values were observed with the co-assembly for all types of environments (Fig 3A) and for each metagenome with the exception of reservoir sediment MG10 (Fig. S4, Table 3). The application of normalization methods (as described in the section below) revealed contrasting patterns in *hgcA* relative abundance, with higher values observed for single assembly methods when applying, for instance, a normalization method based on *rpoB* coverage values (Fig. S4). For each metagenome, the nMDS analysis showed a high level of similarity in taxonomy-based *hgcA* inventories obtained from single assembly vs co-assembly (Fig. 3B). This was confirmed by a procrustean analysis that showed significant levels of concordance for the *hgcA* inventories obtained between both approaches (p ≤ 0.001). Looking at each dataset independently, reservoir sediments and brackish waters showed significant levels of concordances (p ≤ 0.008, p ≤ 0.002) while lake waters and paddy soils had non-significant levels of concordances (p ≤ 0.17, p ≤ 0.30; no statistics possible with only two metagenomes for lake sediments).

**Figure 3.**
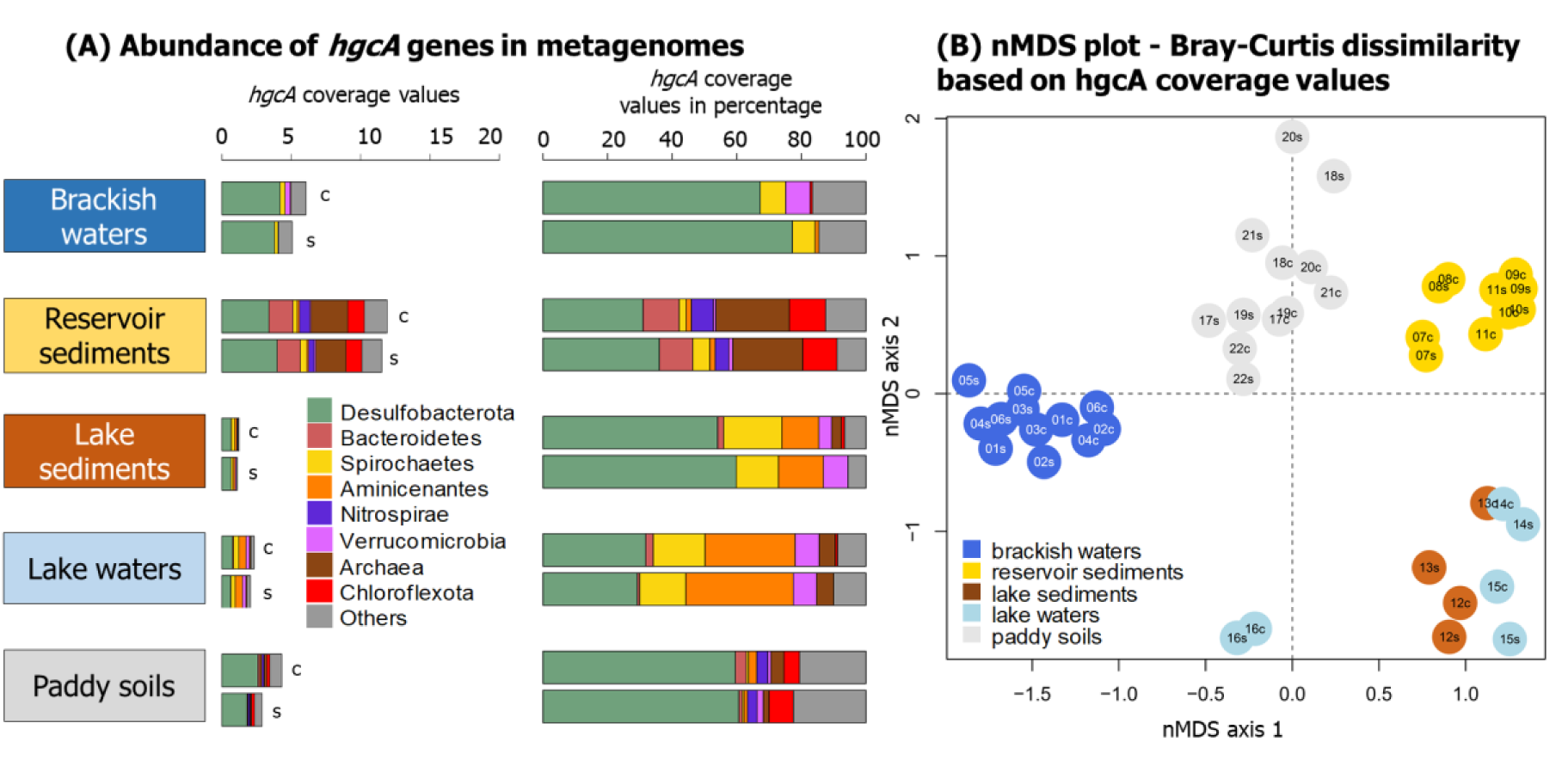
(A) Distribution of *hgcA* genes in the metagenomes obtained from five types of environments with the co-assembly ‘c’ and the single assembly ‘s’ methods. For these barplots, unnormalized hgcA coverage values were used. (B) Dissimilarities in the structure of *hgcA* inventories obtained with the co-assembly ‘c’ and the single assembly ‘s’ approaches. nMDS stress values = 0.1909. The id of each metagenome is denoted as follows: numbers corresponding to the metagenome id (e.g., MG01 is 01), ‘c’ or ‘s’ stands for analysis with the co-assembly or the single assembly.

### 3.4 Comparison between normalization methods

In order to compare normalization methods to estimate the abundance of *hgcA* genes, we calculated the (i) total number mapped prokaryotic reads, (ii) *rpoB* genes coverage values, (iii) median coverage value of 257 marker genes and (iv) genome equivalents values (Microbe Census) (Fig 4, Datasheet 1E). Overall, significant correlations were observed between the total number of reads, *rpoB* coverage values, and the median coverage values of 257 marker genes (Fig. 4A), while no significant correlations were observed between these metrics and genome equivalent values. The 3D plot shows the relationships between the total number of reads, the median coverage values of 257 marker genes and genome equivalent values (Fig. 4B).

**Figure 4.**
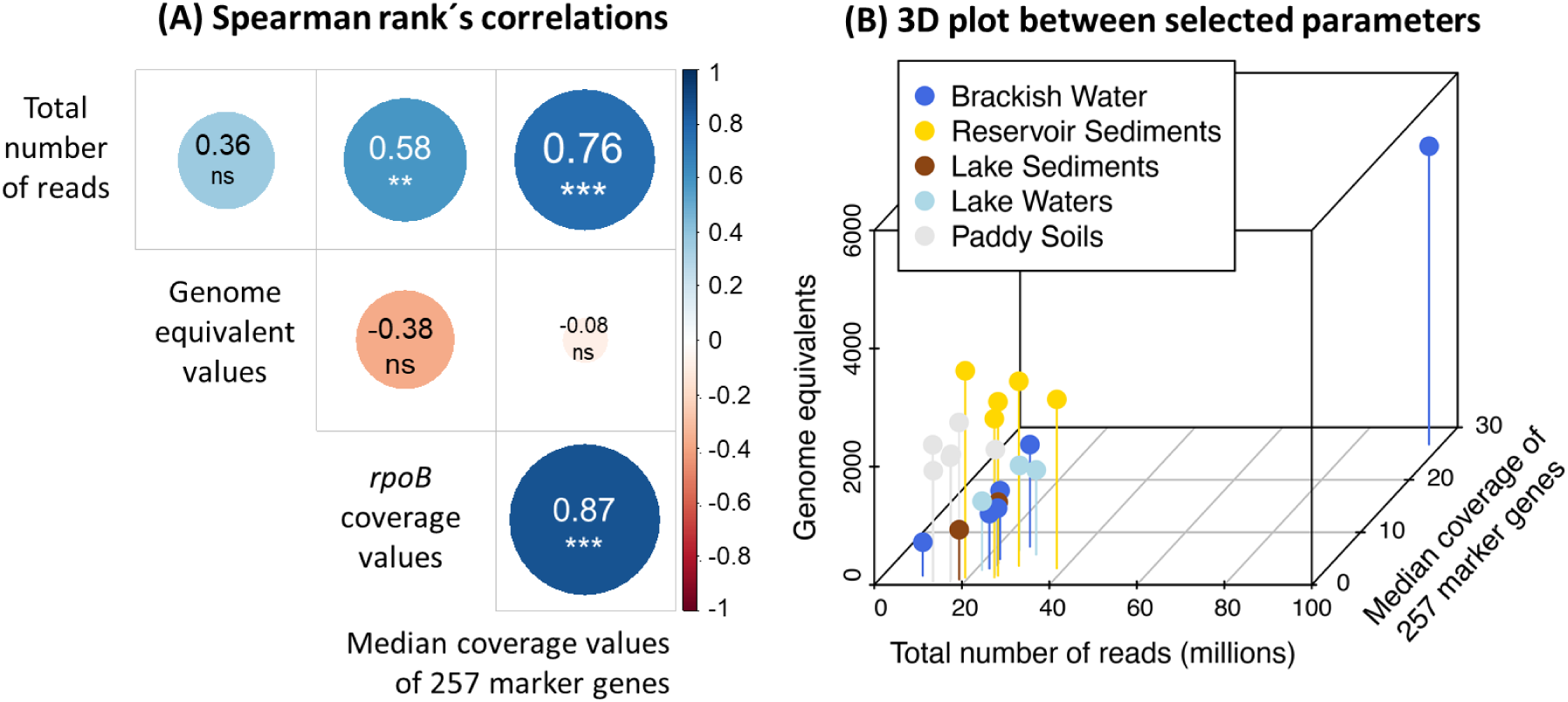
Plots showing correlations between metrics used for normalization. Only outputs presented here were calculated from data obtained with the single assembly approach.

## 4. Discussion

### 4.1 Identification of true *hgc* genes from environmental genomic data

The absence of cultured representatives of *hgc*^+^ microorganisms from novel clades (i.e., outside the Desulfobacterota, Firmicutes, Methanomicrobia) with experimentally validated Hg-methylating capability (Gilmour et al., 2013; 2018) hampers confirmation that newly discovered *hgc* genes from environmental samples truly code for Hg methylating enzymes. Indeed, the recent analysis of publicly available metagenomes revealed the high diversity of microbial lineages with *hgc*^+^ microorganisms, with the vast majority yet uncultured and therefore unstudied for Hg methylation activity (Gionfriddo et al., 2019; McDaniel et al., 2020). To date, all *hgcA*^+^ microorganisms that have been experimentally tested have been found shown to produce MeHg (except for those with fused *hgcAB*-like sequences) (Gilmour et al., 2013; 2018), and protein modeling of novel *hgcA* sequences suggest they have comparable active sites to HgcA sequences in experimentally verified Hg methylators. Therefore, although recent findings revealed relationships between microbial expression of *hgc* transcripts and MeHg formation in the environment (Capo, Feng et al. 2022 bioRxiv), and some putative *hgcAB* genes have been computationally modelled to possess functionality for methylation (Gionfriddo et al. 2016, Lin et al. 2021), we remain cautious about defining true *hgc* genes from environmental samples. As such, some studies have qualified *hgc* genes found in the environment as *hgc* genes (e.g., Gionfriddo et al., 2016; Bowman et al., 2020; Villar et al., 2020; Capo et al., 2020).

Here, we defined three stringency cutoffs to describe *hgcA* genes in environmental metagenomes. By definition, the HgcA-encoding genes detected with the high stringency cutoffs include the key amino acid residues (i.e., the cap helix motif N[V/I]WC[A/S][A/G/S]GK), Parks et al., 2013) present in HgcA from known Hg methylators. In contrast, all other hits to the HMM, from moderate and low stringency cut-offs, lack these amino acid residues. To date none of the isolates lacking these key amino acid residues has been found to methylate Hg, or no cultured isolate exists to test for Hg methylation capability (Gilmour et al., 2018). Substitution of some of these amino acids in the cap helix of HgcA may not result in loss of Hg methylation activity, as demonstrated by site-directed mutagenesis experiments with *P. mercurii* ND132 (Smith et al., 2015). However, in addition to the cap helix domain of HgcA, the transmembrane domain of HgcA may also be required for Hg methylation activity. Unfortunately, the transmembrane region of HgcA has no detectable sequence homology (Cooper et al., 2020).

Thus, we recommend using the high stringency cutoff defined in the present study for routine identification of *hgcA* from environmental metagenomes. Lower stringency could reveal novel HgcA sequences that have lower similarity to HgcA from known Hg methylators, but if the lower stringency cutoff is used, we advise careful manual inspection of the sequences to ensure that they have important motifs and other HgcA features like the cap-helix region. If the amino acid sequence in the cap helix domain is highly divergent from known sequences, we recommend protein modeling efforts to determine if the active site is similar enough to known sequences to validate classification as HgcA. Additional verification of true HgcA sequences include prediction of transmembrane domain regions (e.g., using TMHMM software, Krogh et al., 2001) and identification of other key conserved residues (Parks et al., 2013; Smith et al., 2015; Jones et al., 2019). A combination of several methods will certainly help to improve our description of *hgcA* genes in the coming years.

### 4.2 Effectiveness of the Hg-MATE database

The Hg-MATE database originates from the combination of two recent works (Gionfriddo et al., 2019; McDaniel et al., 2020). The present work is a collaborative project of the Meta-Hg working group that aimed to provide a living database that will be periodically updated. It provides several useful tools (HMM profiles and references phylogenetic trees) and a documented workflow that allows for the identification of *hgc* genes for easy comparison between studies. One major advantage of Hg-MATE is the assignment of NCBI taxonomy IDs (txid) to *hgcA* genes allowing for easy comparison with datasets from other studies that also use the Hg-MATE database (Datasheet 1B). In contrast, outputs from previous *hgc*-related studies are difficult to compare with each other because *hgc* taxonomic identification is usually done with different in-house databases and/or phylogenetic tools, and is based on the manual inspection of phylogenetic trees increasing the level of uncertainties and subjectivity in taxonomic identification. While the used *pplacer* approach here is not perfect - since phylogenetic relatedness of the gene does not necessarily mean the same organismal taxonomy because of potential horizontal gene transfer (McDaniel et al., 2020) - it is a standardized approach allowing for a robust and automated identification of *hgc* genes from metagenomes.

A side-by-side comparison of previous and present taxonomic identification of putative Hg methylators is presented in this section. For water and sediment metagenomes from Lake Manganika our identification by HgcA phylogeny showed consistent results with previous identification from *hgc*^*+*^ MAGs (Jones et al., 2019), with Desulfobacterota, Aminicenantes, Kiritimatiellaeota and Spirochaetes being the predominant putative Hg methylators. In the case of Baltic Sea water metagenomes, the comparison of our Hg-MATE taxonomy identification with the previous identification using a set of *hgc* sequences from Podar et al. (2015) revealed consistency in the predominant *hgc*^+^ groups detected (Desulfobacterota, Spirochaetes, Kiritimatiellota) but noticeable differences for others i.e., Planctomycetes and Verrucomicrobia (Datasheet 1B). Consistent with previous characterization, reservoir sediments were characterized by predominant *hgc*^+^ Methanomicrobia, Desulfobacterota, Bacteroidetes, and Chloroflexota. Finally, in paddy soils, Liu et al. (2018) identified mostly *hgc*^+^ Desulfobacterota, Firmicutes and Methanomicrobia while, in the present study, the two last microbial groups were found less predominant to the benefit of *hgc*^+^ Nitrospirae and Chloroflexota.

In addition to using phylogenetic placements of *hgc* genes in reference trees from the Hg-MATE database, a more precise approach to identification of putative Hg methylators is probably the identification of *hgc*^*+*^ MAGs (i.e., Jones et al., 2019; Peterson et al., 2020; Lin et al., 2021). However, the recovery of MAGs from metagenomes is not always possible due to (i) the difficulty of obtaining MAGs from certain environments such as sediments and (ii) the low predominance of Hg methylators compared to other microorganisms in the environment, and therefore the lower probability of recovering *hgc*^+^ MAGs. A recent work revealed the good congruence between the identification of hgc^+^ MAGs and a *hgc* phylogeny based on Hg-MATE phylogeny (Capo, Feng et al., 2022 bioRxiv) highlighting that both approaches could be used to ensure the reliability in the identification of Hg methylators.

### 4.3 Assembly methods depend of the diversity of the metagenome

The increasing amount of publicly available environmental genomic data (Thompson et al., 2017; Nayfach et al., 2021) opens avenues to answer ecological questions related to the biogeography patterns and dispersal barriers of Hg methylators in interconnected systems (such as the global ocean and coastal systems). Co-assembly of multiple metagenomes has been shown to have many important benefits compared to single assemblies including improved binning and better recovery of low abundance environmental genomes from studies that use multiple low-coverage metagenomes. However, co-assembly requires higher computational costs and potentially masks microdiversity by collapsing the genomes of multiple related strains into a single MAG (Narasingarao et al., 2012; Van der Walt et al., 2017; Ramos-Barbero et al., 2019; Tamames et al., 2020; Paoli et al., 2021). Here, we compared *hgcA* recovery from single assembled metagenomes versus co-assemblies of multiple metagenomes from the same environment. In all cases except one, co-assembly significantly increased the recovery of *hgcA* genes (Fig S4). Additionally, we showed that when the diversity and composition of the *hgcA*^+^ community was compared across all the samples included in the analysis, single assemblies and co-assemblies performed similarly in this regard, suggesting that also single metagenomes can provide adequate information (similar level of *hgc* coverage and detected diversity) on the *hgc*^+^ community.

Differences in the diversity of environments can have an effect on the recovery of *hgc* genes from metagenomes. Nonpareil diversity index values of the metagenomes ranged between 18.7 and 23.7 with the highest being found in paddy soils metagenomes (Fig. S1, Datasheet 1A). Here, for the paddy soils that exhibited higher Non-pareil diversity index values (Fig S1), consistently with Rodriguez-R and Konstantinidis (2014), the co-assembly approach outperforms single sample assemblies in the recovery of *hgc* genes (Fig 3). Noticeably, although no identical HgcA amino acid sequences were detected between single assembly and co-assembly approach, identical DsrA amino acid sequences were observed. We hypothesized that the low proportion of *hgcA* genes in metagenomes, compared to *dsrA* genes, explained such discrepancies, although it did not strongly impact the overall *hgcA* coverage values recovery. In these situations, we recommend aiming for either higher depth of coverage or sequencing of multiple adjacent or linked metagenomes or replicates from a single sample. In contrast, we recommend avoiding the co-assembly of metagenomes from different environments that could produce more misassembles and chimerism (Mikheenko et al., 2016; Sczyrba et al., 2017; Tamames et al., 2020). For other environments such as brackish and lake waters, our work highlights that using the marky-coco pipeline based on a single assembly approach provide similar results to a co-assembly approach in detecting *hgc* genes.

### 4.4 Robust normalization methods are needed for quantitative inferences

The normalization of gene counts from environmental metagenomes and metatranscriptomes is a key aspect of works aiming to study the prevalence of certain microorganisms in specific environments (Pereira et al., 2018; Salazar et al., 2019; Pierella Karlusich et al., 2022). In *hgcAB* omics studies, the number of mapped reads and the coverage values of marker genes or housekeeping genes is usually used to normalize the coverage values of *hgc* genes (Lin et al., 2021; Vigneron et al., 2021; Tada et al., 2021; Capo et al., 2022). Tests here revealed that a wide range of contrasting normalization methods all provided reasonable abundance estimates that were significantly correlated with one another with the exception of genome equivalent values (Fig 4). Non-significant correlations found between genome equivalent values and other metrics can be explained by the weaker relationships observed for the metrics in paddy soils and reservoir sediments metagenomes, while metrics from brackish waters, lake sediment and waters appear to have linear relationships. Therefore, we do not strongly recommend any single method over others. Instead, we suggest that it may be prudent to report data that employ multiple normalization methods to allow for easy comparisons to be carried out between studies. Such normalizations can without too much of an effort be included in the supporting information for later usage. Suggested normalization methods include the total number of prokaryotic reads, coverage values of *rpoB* genes and the median coverage values of 257 marker genes (example in Datasheet 1E).

## 5. Conclusion

The study of the taxonomic diversity and metabolic capacities of microorganisms involved in Hg methylation will lead to a better understanding of the environmental factors triggering microbial methylation of divalent Hg. Although metagenomic and metatranscriptomic-based studies have provided better insights into the environmental role of those microorganisms, there is still a need to standardize methods to detect *hgc* genes from environmental omic data. Furthermore, since Hg methylators often constitute such a small proportion of the microbiome, methods outlined in this study provide best practices for improving their detection and recovery from metagenomes. We provide here an up-to-date *hgc* gene catalogue, Hg-MATE database v1, and the marky-coco bioinformatic pipeline to detect, identify and count *hgc* genes from metagenomes. We recommend using our high stringency cutoff to detect hgcA genes in metagenomes and applying our protocol in future prospects of Hg methylation genes, especially for cross-comparison between studies. Finally, although a co-assembly approach should be chosen when analyzing metagenomes from highly diverse environments (i.e., paddy soils), we recommend using marky-coco pipeline, based on a de novo assembly for recovering *hgc* genes in metagenomes from aquatic environments.

## Acknowledgements

This work was funded by the Severo Ochoa Excellence Program postdoctoral fellowship awarded in 2021 to Eric Capo (CEX2019-000928-S), the Swedish research council Formas (grant 2018-01031), the EMFF-Blue Economy project MER-CLUB (grant 863584). Caitlin Gionfriddo was a Robert and Arlene Kogod Secretarial Scholar with the Smithsonian Environmental Research Center while conducting work described in this manuscript. Benjamin Peterson was funded as a postdoctoral research associate by the National Science Foundation (Award 1935173) during the work in this manuscript. The computations were performed on resources provided by SNIC through Uppsala Multidisciplinary Center for Advanced Computational Science (UPPMAX) using the compute project SNIC 2021/5-53. Some of the computations for compiling the Hg-MATE database were conducted on the Smithsonian High Performance Cluster (SI/HPC), Smithsonian Institution. https://doi.org/10.25572/SIHPC. The authors are thankful to the members of the Meta-Hg working group (https://ercapo.wixsite.com/meta-hg) and Mersorcium network (https://mersorcium.github.io/).

## Conflict of interest

The author declares no conflict of interest

## Benefit-sharing statement

Benefits from this research is the creation and curation of Hg-MATE database (https://doi.org/10.25573/serc.13105370.v1) and release of the bioinformatic pipeline marky-coco (https://github.com/ericcapo/marky-coco).

## Data availability statements

All metagenomes analyzed in this study are of public access as described in Table 2.

**Datasheet 1**. This file includes information related to different parameters collected or measured in this work from the 29 metagenomes used in this work (A) For each metagenome, metagenome id, type of environment, non-pareil metrics, genome equivalents (Microbe Census) values, number of cleaned and mapped reads, number of *hgcA* genes, *hgcA* coverage values and normalization metrics values, *dsrA* coverage values (B) List of all *hgcA* genes detected in the 29 metagenomes with both a single assembly and co-assembly approaches, with the three stringency cutoffs. Gene length, number of mapped reads, coverage values, NBCI taxonomy txid and amino acid sequences are presented. (C) List of all *hgcA* genes detected in the 29 metagenomes with both a single assembly and co-assembly approaches, with the high stringency cutoff. Gene length, number of mapped reads, coverage values, NBCI taxonomy txid and amino acid sequences are presented. (D) List of all *dsrA* genes detected in the 29 metagenomes with both a single assembly and co-assembly approaches, with the high stringency cutoff. Gene length, number of mapped reads, coverage values and amino acid sequences are presented. (F) Coverage values of the 257 marker genes (including *rpoB*) obtained using the single assembly vs co-assembly approaches.

## Supporting Information

**Figure S1:**
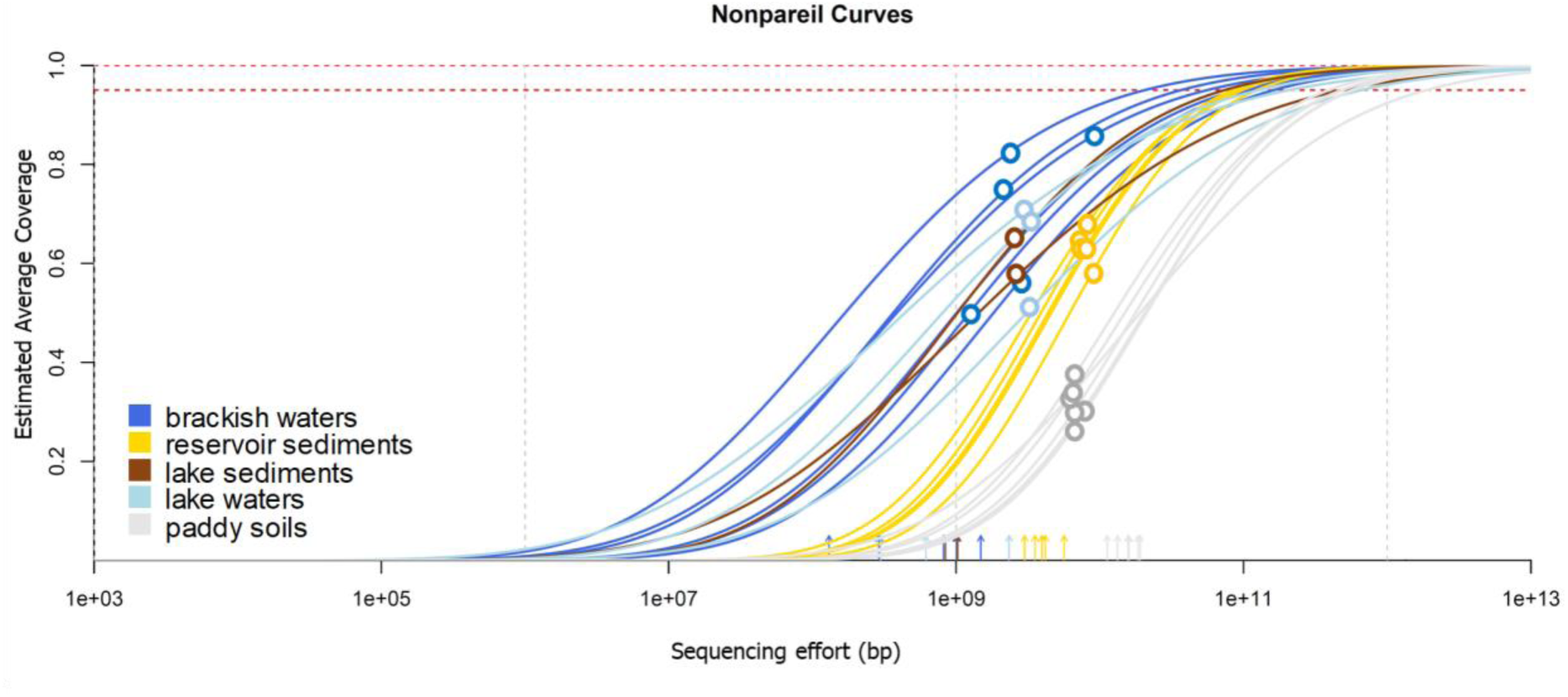
Nonpareil curves for the 22 metagenomes. The plot displays the fitted models of the Nonpareil curves. The horizontal dashed lines indicate 100 (gray) and 95% (red) coverage. The empty circles indicate the size and estimated average coverage of the datasets, and the lines after that point are projections of the fitted model.

**Figure S2.**
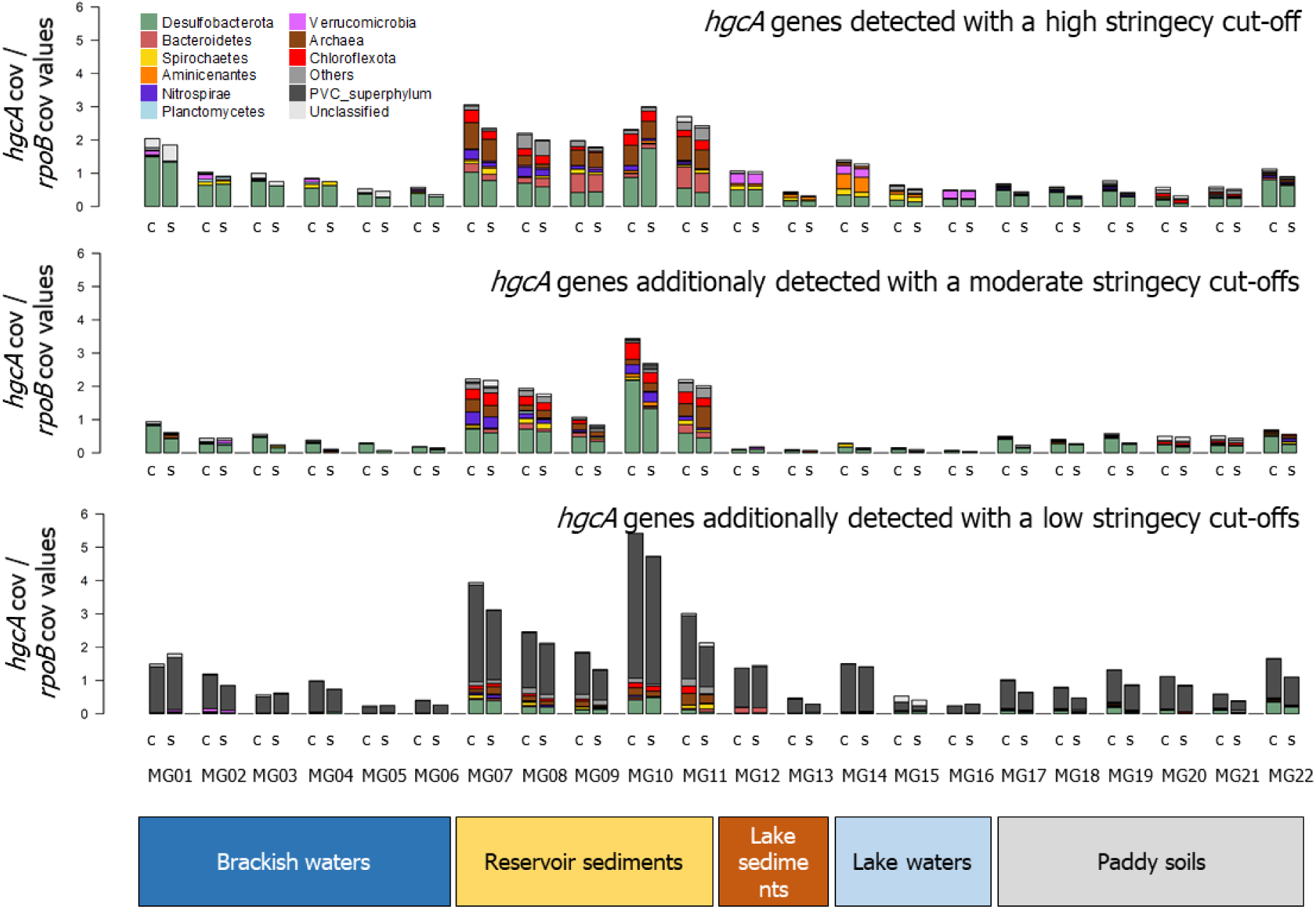
Distribution of *hgcA* genes in the 22 metagenomes recovered using the co-assembly ‘c’ and the single assembly ‘s’ methods and applying the three stringency cutoffs defined in this manuscript for the definition of *hgcA* genes. Abundance values were calculated as *hgcA* coverage values normalized by *rpoB* normalized values. Colors denote taxonomic affiliations of *hgcA* genes.

**Figure S3:**
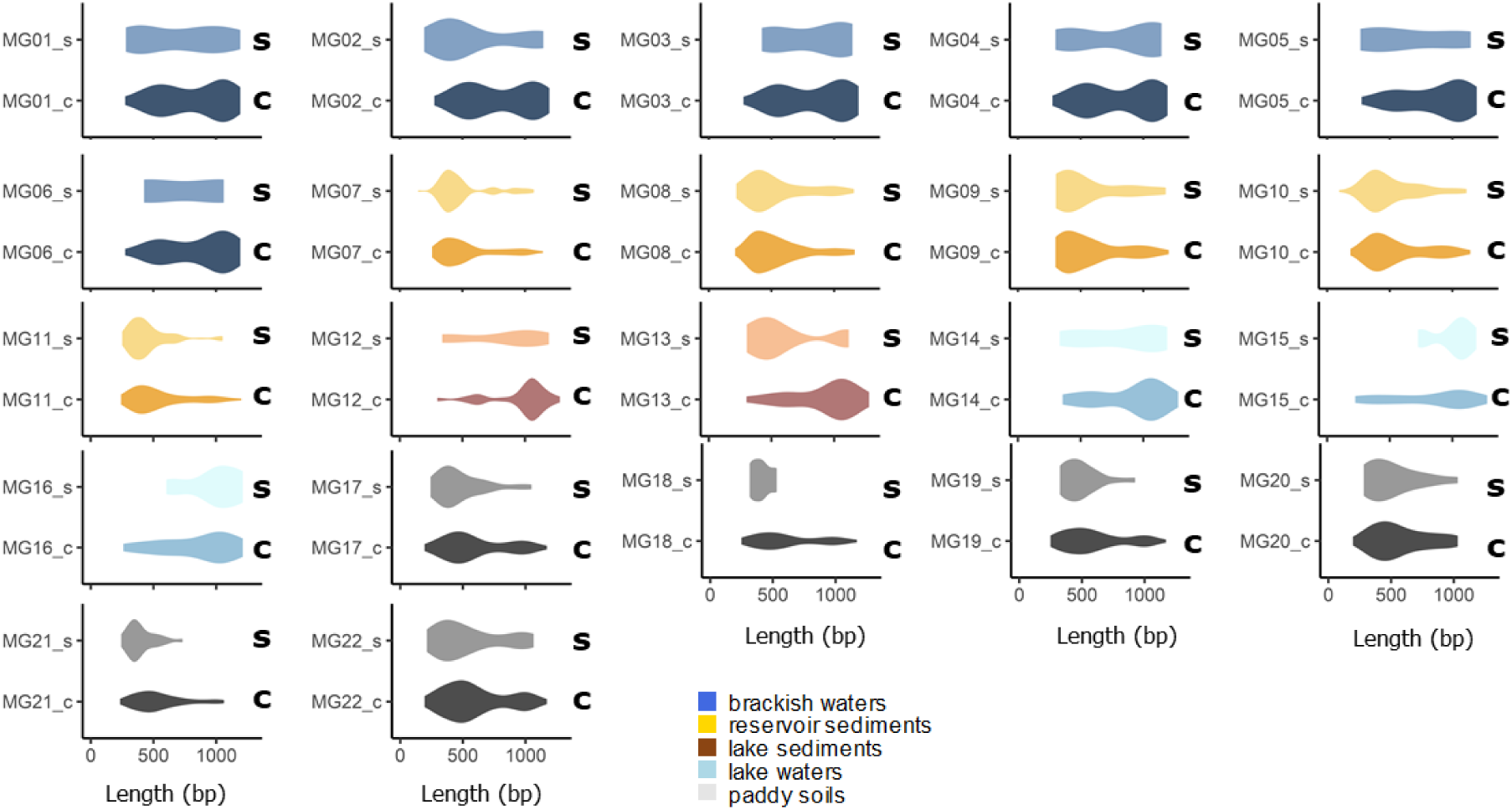
Violin boxplots showing, for each metagenome, the difference in *hgcA* sequence length distribution comparing the outputs of the co-assembly and the single assembly approaches.

**Figure S4:**
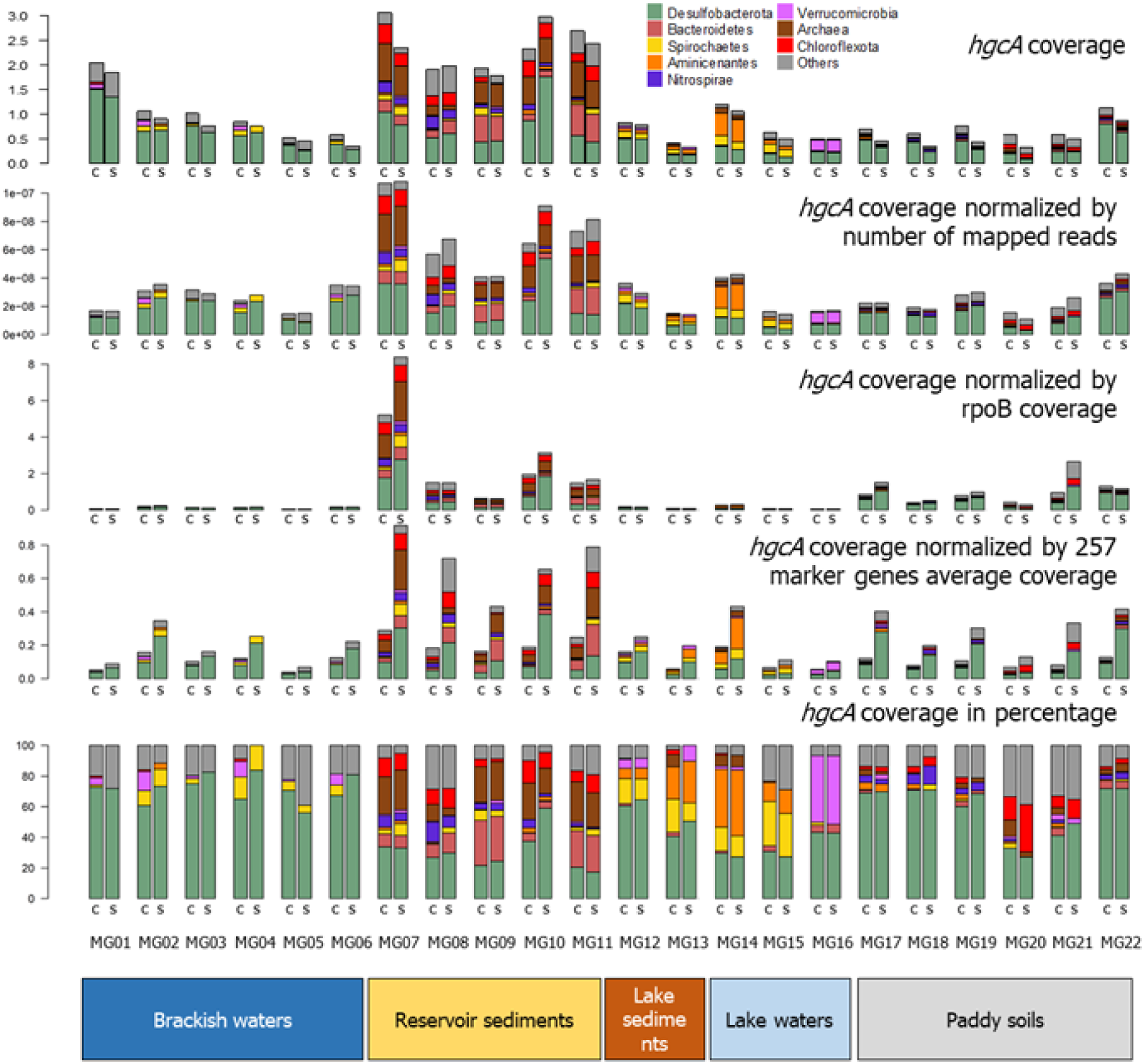
Distribution of *hgcA* genes in the 22 metagenomes with the co-assembly (c) and the single assembly (s) methods with different normalization methods

## Notes

https://github.com/ericcapo/marky-coco

https://smithsonian.figshare.com/articles/dataset/Hg-MATE-Db_v1_01142021/13105370/1?file=26193689

### Competing Interest Statement

The authors have declared no competing interest.

